# Building Key Populations HIV Cascades in Data-Scarce Environments: Towards a participatory stakeholder methodology for cascades construction, adoption, and utilization

**DOI:** 10.1101/452417

**Authors:** Tim Lane, Mike Grasso, Andrew Scheibe, Grace Liu, Alexander Marr, Pelagia Murangandi, Getahun Aynalem, Mariette Slabbert, Lebowa Malaka, Zachary Isdahl, Thomas Osmand, Patrick Nadol

## Abstract

**Introduction:** Recent HIV key populations (KP) surveillance studies in South Africa, including female sex workers (FSW) and men who have sex with men (MSM), demonstrate the disproportionate burden of HIV they bear compared to the general population. The national response for KP has lagged due to relatively scarce KP data focused narrowly on urban areas. We adopted a participatory data triangulation approach with stakeholders to overcome the challenges of KP program planning in KP data-scarce districts. Here we describe our methodology for achieving consensus on population size estimates (PSE) and treatment cascade indicator estimates derived from FSW and MSM surveillance data and applied across the country.

**Methods:** The South African National AIDS Council (SANAC) convened the group; technical advisors from the University of California San Francisco (UCSF) facilitated; and stakeholders from government, non-government, academic, and KP-led advocacy organizations involved in program implementation and research provided input through three in-person meetings covering four phases of work: surveillance data analysis; cascades data extraction; presentation for feedback; and data extrapolation.

**Results:** Technical advisors presented eight cascades (three FSW, five MSM) to stakeholders, recommending data-informed extrapolation factors for each population. Stakeholders adopted recommendations by consensus with few adjustments. FSW cascades displayed high awareness of HIV status and steep breakpoints towards ART uptake; MSM cascades displayed less HIV status awareness, but relatively good ART uptake, with metropolitan areas displaying better uptake than rural districts.

**Conclusion:** The participatory process enabled KP stakeholders to vet disparate data sources against programmatic experience and recommend consistency in cascades data; participatory triangulation of additional surveillance and program data will follow. The considerable time and resource investments in this process had downstream benefits, including consistency in sub-national HIV implementation plans. We recommend this consensus-based approach as a transparent, consistent, and sound methodology for cascades construction in KP data-scarce environments.

## Introduction

Key populations (KP), including female sex workers (FSW), men who have sex with men (MSM), people who inject drugs (PWID), and transgender women are marginalized, stigmatized, and often criminalized throughout Sub-Saharan Africa, and bear a disproportionate burden of HIV(1-4). Despite its relatively enabling human rights-based policy environment, South Africa is no exception to regional trends. Its 1996 constitution decriminalized homosexuality, but MSM and other sexual minority populations remain highly stigmatized; sex work and the use of drugs remains illegal(5). Recent KP HIV surveillance studies in South Africa have demonstrated consistently high HIV prevalence among KP groups—for example, among the country’s estimated 150 000 sex workers(6) prevalence estimates range from 40-88%(7, 8); as many as a third of the country’s 1.2 million MSM(9) are HIV-infected(10); among the estimated 67,000 PWID(11), HIV prevalence ranges from 14-16.2%(5, 8). And, despite significant progress in increasing access to anti-retroviral therapy (ART) across most population groups, including the adoption of Universal Test and Treat (UTT) guidelines (12, 13) in 2016, available evidence suggests significant barriers to treatment that less than one-third of HIV-positive FSW and less than one-quarter of HIV-positive MSM were taking ART in 2013 (14, 15).

South Africa’s relatively enabling policy environment for KP is characterized by meaningful FSW and MSM advocacy within civil society and national HIV planning structures(12, 13): meaningful engagement with organizations representing PWID and transgender women has been more limited. South African civil society organizations are represented on the South African National AIDS Council (SANAC); constituent KP “sectors” have guided sponsors including U.S. President’s Emergency Plan for AIDS Relief (PEPFAR), the Global Fund to Fight AIDS, TB and Malaria, and other funding agencies, to progressively increase investment in KP-focused programs. To date most programs have had limited, generally metropolitan geographic reach, focused on primary prevention strategies and HIV testing services (HTS) promotion. The current National Strategic Plan for HIV, TB and STI’s 2017-2022 has explicitly called for scale-up of comprehensive prevention and treatment programming for KP to smaller urban centers and other areas of identified need (16, 17). Programmatic guidelines for FSW and MSM are laid out in the South African National Sex Worker HIV Plan, 2016-2019(12) and the South African National LGBTI HIV Plan, 2017-2022(13), respectively. These were developed against the backdrop of the 2014 UNAIDS 90-90-90 targets, i.e. 90% of all people living with HIV (PLHIV) know their HIV status, 90% of these on ART (81% of PLHIV), and 90% of these (73% of PLHIV) be virally suppressed(18-20) by 2020. Achieving 90-90-90 targets for South Africa’s KP requires a data-driven approach, prioritizing evidence-based interventions to ensure that KP flow efficiently, consistently, and sustainably through the continua of HIV prevention and treatment services. Thus there is increased need to build and utilize data systems that effectively monitor the care continuum, or cascade, for KP (21).

Cascade analyses provide a framework for assessing and improving service delivery at each stage of HIV care. They are also a logical modeling tool to identify gaps (referred to as “breakpoints” or “drop-offs”) and opportunities for KP-specific interventions in the continuum of HIV, allowing program implementers at facility, regional or national levels to target resources and interventions more effectively, ultimately improving PLHIV engagement in care, viral suppression, and prevention of onward transmission.(22) Yet, the same structural factors that contribute to the inequitable burden of HIV among KP and sub-optimal retention in HIV treatment services (17, 23, 24), also contribute to the scarcity of KP-specific strategic information, and the concentration of what exists in metropolitan health services environments. Until a wider range of sub-national data exists, a transparent, consistent, and methodologically sound approach to triangulating existing data to diverse sub-national urban, peri-urban, and rural environments is critical for proportional HIV program planning(25).

Here, we describe our experience of utilizing data from eight South African KP surveillance sites (three FSW and five MSM) to construct treatment cascades for all 52 sub-national districts for these two populations. We present indicators for surveillance sites as cascades, and describe the methods and results of the stakeholder consensus process we believe generated the best possible data for programmatic planning at this time. We conclude with brief recommendations for roles and responsibilities of conveners, technical advisors, and stakeholders to aid national-level KP HIV programs confronting the need for evidence-based service delivery targets in data-scarce environments.

## Methods

We adapted a method described by WHO for cascade analysis as a group consensus activity involving multiple stakeholders and disparate sources of data. In our case, we relied primarily on IBBS data and stakeholder opinion and experience to arrive at consensus cascades.(26) Our national KP stakeholder consensus process was convened by SANAC; facilitated by technical advisors (TAs) from the University of California San Francisco (UCSF); and stakeholders representing government, non-government, academic, and KP-led advocacy organizations involved in program implementation and research. The work proceeded in four phases. In Phase 1, TAs calculated population size estimates (PSE) for each population, triangulated from multiple PSE methods embedded in integrated bio-behavioral surveillance (IBBS) surveys(27). In Phase 2, the technical advisors used IBBS-derived indicator estimates and PSEs to construct treatment cascades. In Phase 3, investigators presented the methods and results of Phases 1 and 2 to a group of national KP service providers and stakeholders convened by SANAC; facilitated stakeholder discussion on the assumptions, methods, results, and limitations of IBBS data; sought consensus either to adopt estimates as presented, or revise assumptions and calculations for surveillance data-derived cascades for each population and site; and sought consensus on a set of “reasonable” assumptions and factors for PSEs and treatment cascade indicators to apply uniformly throughout the country. In Phase 4, TAs constructed a final set of cascades for the country’s 52 districts, populated according to the consensus outcomes of Phase 3, and presented as a report to SANAC and the National Department of Health (NDOH). Below we reference relevant sampling, survey, laboratory, and PSE methods that informed Phases 1 and 2, and relevant consensus processes that informed Phases 3 and 4.

### Phase 1: IBBS survey and PSE analysis

IBBS data sources were the South Africa Health Monitoring Survey for Female Sex Workers (SAHMS-FSW), fielded at 3 metropolitan district sites (Johannesburg, Cape Town, and eThekwini/Durban) in 2013-14, and the South Africa Men’s Health Monitoring Survey (SAMHMS) for MSM, fielded at 3 metropolitan (Johannesburg, Cape Town, Mangaung/Bloemfontein) and 2 district municipality (DM) sites (Capricorn DM/Polokwane, Limpopo provincial capital; and NM Molema DM/Mahikeng, NorthWest provincial capital) in 2015-16. In each survey, we followed standard second-generation surveillance guidelines for IBBS, including respondent-driven sampling (RDS) survey methodology, that have been described elsewhere(15). Indicator definitions are summarized in Figure 1. Below, we briefly review each IBBS’s laboratory and statistical analysis methods, and describe methods for calculating preliminary population size estimates from IBBS data.

**Figure 1.**
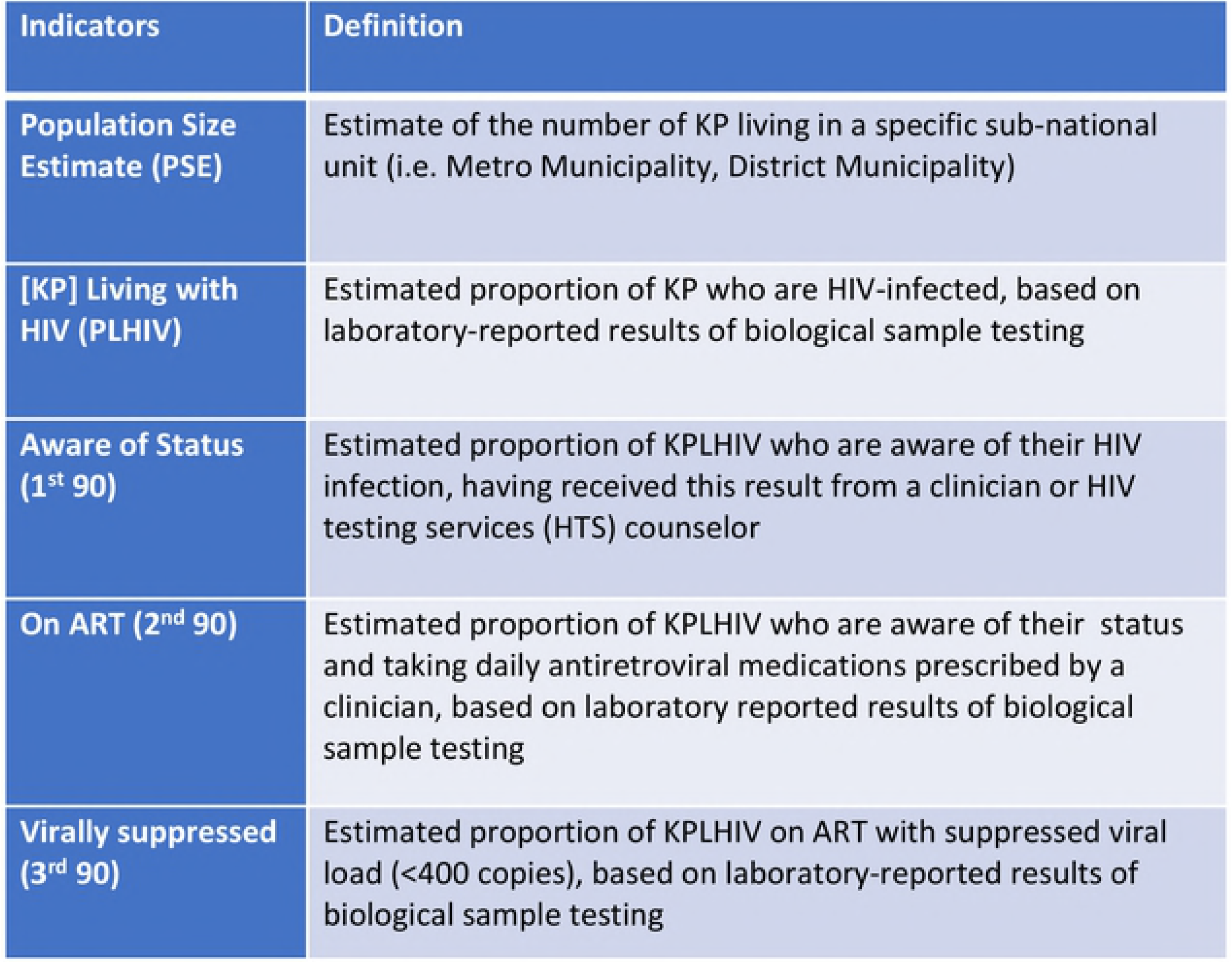
South Africa Key Populations Treatment Cascade Indicator Definitions.

#### Laboratory analysis

We assessed HIV-status by laboratory analysis in both populations. Laboratory methods for SAHMS-FSW are described elsewhere.(7) Briefly, BARC laboratories (Johannesburg) performed testing on serum using an HIV ELISA 4th generation; with confirmation of discordant results using a third-generation HIV ELISA assay. HIV-1 and HIV-2 Western blot (Bio-Rad Laboratories, USA) were run on samples when required for definitive diagnosis.

For MSM, dried blood spot (DBS) cards (Whatman 903 Protein Saver Cards; Sigma-Aldrich, St Louis, MO) were assayed at the National Institute for Communicable Diseases (Johannesburg), screened with a 3rd generation HIV ELISA (Genscreen HIV-1/2 Version 2, Bio-Rad, Marnes-la-Coquette, France). Non-reactive tests were interpreted negative; reactives were confirmed positive by a 4th Generation ELISA, (Vironostika, HIV Ag/Ab Assay, bioMérieux, Marcy-l’Etoile, France), with HIV-1 Western Blots for final HIV infection status. We tested DBS samples for ART analytes at the Division of Clinical Pharmacology, Department of Medicine, University of Cape Town with High Performance Liquid Chromatography coupled to Tandem Mass Spectrometry. Qualitative detection of Nevirapine, Efavirenz, or Lopinavir, was carried out by a validated method using minor modifications of the method used by Koal et al.(28).

#### Statistical analysis

Data from each survey were originally analyzed and presented as RDS-adjusted proportions and 95% Confidence Intervals (CI) using RDS Analysis Tool v.7(4). For FSW, treatment cascade indicators were reported as RDS-adjusted proportions. For MSM, small cell sizes for ART indicators at all sites except Johannesburg returned illogical RDS-adjusted 95% CI values; thus we present here all cascades for all MSM sites as unadjusted sample proportions and 95% CIs.

#### Population size estimation

We calculated PSEs for each survey location for each population as the median of a plausible range of the several methods of PSE calculation described below, setting upper and lower plausibility boundaries. The several methods for each population included IBBS-derived multipliers and modified Delphi (“Wisdom of the Crowds”), and proportional estimates derived from academic literature.

For multipliers, we followed the “multiple multipliers” recommendations of Abdul-Qader and colleagues(27) including a unique object and event multipliers, and service multipliers where available. Point estimates were derived according to the formula *N=n/p* where *N* is the population size; *n* the number of objects (e.g. bracelets) distributed in the target population, or attendees at a memorable event (e.g. survey launch party), or number accessing population-specific services (e.g. HTS) over a defined time period prior to the survey; and *p* is the proportion of survey participants self-reporting receipt of an object, attendance at the event, or utilization of the service, respectively. Additionally, we calculated the modified Delphi indicator (“Wisdom of the Crowds”) as the mean of two responses to “How many [target population] do you think there are in and around [surveillance site city/district]?” posed at the beginning and conclusion of each survey; and the PSE for the site as the mean of the middle 90^th^ percentile of all responses.

We then used literature-derived estimates to explore whether our preliminary median and plausible range results were consistent with other empirical data and expert opinion from South Africa and elsewhere. We compared recommended proportions to the 2011 census-derived adult population of females (FSW) and males (MSM) aged 15-49 (the approved protocol for each survey permitted inclusion of minors at least 16 years of age; for FSW, per protocol, individuals under age 18 underwent additional screening for evidence of trafficking and were referred to appropriate social services), based on the degree of urbanicity (large metropolitan municipalities versus smaller provincial district municipalities). For FSW, this range was 0.4-0.6% of females described by Konstant and colleagues;(29) for MSM, 1.2%-2.0% described by Caceres and colleagues.(30)

### Phase 2: Data extraction for preliminary cascade analysis

Technical advisors used cascade indicator definitions described by the WHO cascade analysis guide and tool (2017)(26), modified for surveillance analysis by retaining PLHIV as the cascade denominator so that results would demonstrate unmet need towards UNAIDS 90-90-90 targets. Relevant treatment indicator proportions and PSEs were extracted from surveillance data and entered into a Microsoft Excel worksheet to construct three FSW and five MSM preliminary treatment cascades, using the IBBS-based median PSE and HIV prevalence estimates to define the PLHIV denominator, and the subsequent numerator for each indicator calculated as the survey-estimated proportion of PLHIV aware of status and taking ART at each site. (See Phase 3 for viral suppression indicator extrapolation).

### Phase 3: Stakeholder presentation and modified Delphi consensus

Technical advisors (TAs) invited input on preliminary cascades following a Delphi consensus methodology described by the San Francisco (USA) Department of Public Health(31-33) (SFDPH) and previously implemented in Tanzania(34) and Ghana(35) to adopt sub-national key populations size estimates. SANAC convened three stakeholder workshops between September 2016 and February 2018 facilitated by UCSF TAs, who led stakeholders in exploring the surveillance data informing the cascades. At the first face-to-face meeting, investigators explained the intended purpose of the consensus process as involving two steps: the presentation, discussion, and adoption of the surveillance cascade methods and results for each of the FSW and MSM surveillance sites; and the adoption of transparent, consistent, and methodologically sound extrapolation factors from which to construct sub-national cascades for each population in the districts for which no surveillance data existed (49 and 47, respectively). At the second face-to-face meeting, in February 2017, investigators presented preliminary extrapolation results for 11 districts, including 3 of the country’s 4 largest metropolitan areas with over 1.0 million total population, and 7 of 9 provincial capitals of 1.0 million or fewer residents. Stakeholders and investigators assessed the reasonableness of preliminary extrapolation results together, recommended reasonable adjustments, and adopted a final set of extrapolation assumptions and factors for sub-national PSEs and treatment cascade indicators.

Because neither FSW nor MSM had laboratory validated results for viral suppression available for cascades analysis, stakeholders asked TAs to consult programmatic sources of viral suppression data and recommend reasonable extrapolation assumptions. Technical advisors consulted National Health Laboratory Service viral suppression data collected between October 2015 and September 2016, disaggregated by sex and district. Under the assumption that the clinical response of FSW was as well as women generally and MSM as well as men generally on ART, and using the same urbanicity categories previously agreed upon, we calculated extrapolation factors as the mean proportion of the middle 90^th^ percentile of virally suppressed women and men, and applied these respectively to the surveillance or extrapolated proportions of FSW and MSM on ART per district. A final meeting to achieve consensus on the reasonableness of TA’s assumptions and results was convened in February 2018. TAs offered stakeholders offered the alternative of submitting written comments in advance of the meeting.

### Phase 4: Construction of final sub-national FSW and MSM treatment cascades

Technical advisors applied all consensus extrapolation factors to a pivot table pre-populated with 2011 census data disaggregated by sex and age (females 15 years and older; males 15 years and older) to construct 52 final sub-national cascades for FSW and MSM based on definitions and indicator data described above. Technical advisors presented a summary report on behalf of the entire group to SANAC, NDOH, and constituent stakeholders.

## Results

We constructed 8 treatment cascades (3 FSW, 5 MSM) (see Figure 1) with previously analyzed surveillance data (see Table 1). (For convenience and ease of interpretation, we include extrapolated viral suppression estimates in Table 1; these were a Phase 3 extrapolation outcome in our process). By consensus, stakeholders assumed that overall population size and urbanicity might be representative of differences in availability and uptake of health services by KP, and recommended three categories to vet all surveillance cascade indicators against: metropolitan municipalities with total population sizes greater than 1.0 million [A], metropolitan and district municipalities between 200,000 and 1.0 million [B], and district municipalities less than 200,000 [C].

**Table 1:** South Africa Health Monitoring Survey Results: Population Size Estimates and progress towards 90-90-90 indicator targets, 2013-14 (FSW) and 2015-16 (MSM). *90-90-90 targets are expressed with PLHIV as denominator*

|  | Population Size Estimates |  | Prevalence estimates/PLHIV |  | PLHIV Aware of status (Target: 90%) |  | PLHIV On ART§ (Target: 81%) |  | PLHIV Viral Suppression‡ (Target: 73%) |  |
| --- | --- | --- | --- | --- | --- | --- | --- | --- | --- | --- |
|  | (N) | (%)* | % | (n) | % | (n) | % | n | % | (n) |
| **FSW (2013-14)** |  |  |  |  |  |  |  |  |  |  |
| **February 2018 Consensus** |  |  |  |  |  |  |  |  |  |  |
| Johannesburg Metro | 10,895 | 0.7% | 71.8% | 7,823 | 73.8% | 5,773 | 19.0% | 1,486 | 15.2% | 1,189 |
| Cape Town Metro | 7,500 | 0.7% | 40.0% | 3,000 | 56.7% | 1,701 | 27.7% | 831 | 20.5% | 615 |
| eThekweni Metro (Durban) | 9,323 | 0.7% | 53.5% | 4,988 | 77.0% | 3,841 | 25.6% | 1,277 | 22.2% | 1,107 |
| **MSM (2015-16)** |  |  |  |  |  |  |  |  |  |  |
| **September 2016 Consensus** |  |  |  |  |  |  |  |  |  |  |
| Johannesburg Metro | 37,549 | 2.2% | 43.4% | 16,296 | 55.7% | 9,077 | 43.0% | 7,007 | 32.6% | 5,312 |
| Cape Town Metro | 29,901 | 2.2% | 26.7% | 7,984 | 50.3% | 4,016 | 40.0% | 3,194 | 30.4% | 2,426 |
| Capricorn DM (Polokwane) | 5,270 | 1.4% | 22.3% | 1,175 | 24.7% | 290 | 22.3% | 262 | 15.6% | 183 |
| NM Molema DM (Mahikeng) | 3,779 | 1.4% | 18.2% | 688 | 29.4% | 202 | 29.4% | 202 | 20.6% | 142 |
| Mangaung Metro (Bloemfontein) | 3,655 | 1.4% | 18.1% | 662 | 31.8% | 211 | 36.4% | 241 | 25.5% | 169 |

\*PSE % is the proportion of the adult female (for FSW) or male (for MSM) population $\geq 15$ according to the South African Census, 2011.
§ART for FSW are self-reported proportions by site presented in SAHMS 2013-14 report; and laboratory verified proportions by site for MSM in SAMHMS 2015-16
‡Viral suppression estimates are extrapolated from National Health Laboratory Services summary data, October 2016-September 2017, disaggregated by sex and district population size; factors of 80% for FSW and 72% for MSM in metropolitan districts >1.0 million population; and 70% for MSM in smaller metros & district municipalities adopted by stakeholder consensus, February 2018. Proportions of PLHIV virally suppressed presented here are equivalent to the proportion of each population's PLHIV On ART\*extrapolated proportion (e.g. 80%, 72%, etc.) assumed virally suppressed.

Table 2 presents the summarized results of the participatory process as the adopted extrapolation factors for all districts for which surveillance data was not available. For FSW in Johannesburg, Cape Town, and eThekwini, stakeholders originally adopted the PSEs as presented for each of these three cities as the September 2016 FSW Consensus PSEs. Because these results were similar to those observed by Konstant and colleagues in 2013,(29) stakeholders recommended the adoption of extrapolation factors recommended by Konstant to estimate FSW PSE in all other districts (i.e. 0.4%, 0.5%, and 0.6% of the census population figure for adult females aged 15-49). In the three surveillance cities, awareness of HIV status was high (range 56.7%-77.0%), and showed substantial drop-offs towards ART outcomes. There was agreement among stakeholders that knowledge of HIV status was likely similar in all municipalities regardless of size, and recommended for awareness of status proportion, the median 73.8% from surveillance data as a valid extrapolation factor for all districts. For self-reported ART use, stakeholders considered the age of the data (collected in 2013-14), policy shifts (from treatment eligibility of CD4<350 in 2014 to UTT in 2016), and intensified programmatic activity to recommend to TAs that self-reported ART indicator results be multiplied by a factor of 25% for the 2018 cascades.

**Table 2:** South African Key Populations Stakeholder Consensus Indicator Assumptions for Extrapolated HIV Treatment Cascades, 2018.

|  | FSW |  |  | MSM |  |  |
| --- | --- | --- | --- | --- | --- | --- |
| Indicator | SAHMS proportions: median (range) | Metros >1.0m (A) | Metros <1.0m (B) & District Municipalities (C) | SAMHMS proportions median (range)* | Metros >1.0m (A) | Metros <1.0m (B) & District Municipalities (C) |
| PLHIV | 53% (40%-72%) | PSE*0.53 | PSE*0.53 | 22% (18%-43%) | PSE*0.30 | PSE*0.30 |
| Aware of HIV+ status | 74% (57%-77%) | PLHIV*0.738 | PLHIV*0.738 | 36% (23%-53%) | PLHIV*0.53 | PLHIV*0.27 |
| On ART | 26% (19%-28%) | PLHIV*0.256*1.25 | PLHIV*0.256*1.25 | 26% (15%-32%) | PLHIV*0.33 | PLHIV*0.17 |
| Virally Suppressed | n/a | ART*0.80 | ART*0.752 | n/a | ART*0.72 | ART*0.702 |

\*By consensus of stakeholders, MSM Aware of status and On ART indicator extrapolations were considered as means of Category (A) and (B)&(C) municipalities' surveillance results, not as median of the given range.
§ On ART for FSW are self-reported proportions per site presented in SAHMS 2013-14 report; On ART results for MSM are laboratory reported proportions per site in SAMHMS 2015-16
‡Viral suppression results extrapolated from National Health Laboratory Services summary data, October 2016-September 2017, disaggregated by sex and district population size; factors adopted by stakeholder consensus, February 2018.

At the February 2018 meeting, FSW stakeholders presented the group with systematically deduplicated programmatic data showing program reach in Johannesburg and Cape Town at 150% of the upper plausible limit from the September 2016 consensus (eThekwini reach was consistent with the September 2016 PSE). Technical advisors recommended inclusion of the programmatic data as an additional data point to calculate the median at each of the three surveillance sites and presented these results to the group with three options: reject the new estimates; adopt the new estimates for these cities only; or adopt and adjust all sub-national estimates proportionally. The stakeholders recommended adjusting PSE only for the three cities because there was no programmatic evidence from elsewhere in the country to recommend revisiting the previous consensus for these areas. The (n) for each FSW indicator in Table 1 is derived from the February 2018 FSW Consensus PSEs for these three cities; the extrapolation factors listed in Table 2 were not affected by this adjustment. We present selected extrapolated FSW cascade results for three municipalities in Figure 2.

**Figure 2.**
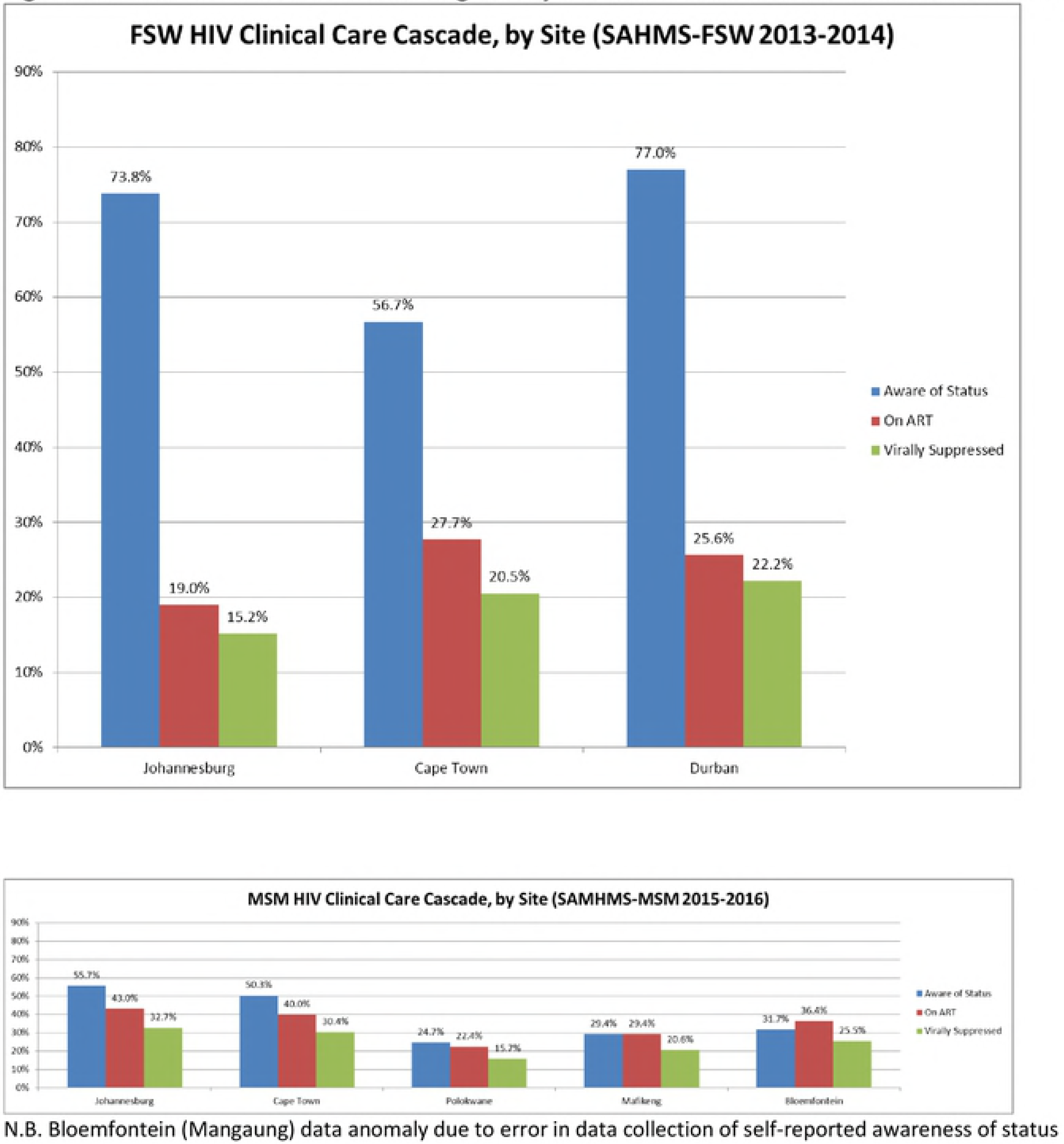
FSW and MSM surveillance survey results displayed as HIV Treatment Cascades.

For MSM, stakeholders adopted the PSEs as presented by investigators as the September 2016 MSM Consensus PSEs, and further adopted the median proportion of 2.2% as an extrapolated PSE for the largest metropolitan areas [A], and the median of 1.4% for all other areas [B and C]. For cascade indicators, MSM stakeholders agreed that a triangulated surveillance and literature-based HIV prevalence of 30% was a reasonable extrapolation assumption across all districts. Noting clustering of cascade curves by urbanicity (i.e. Johannesburg and Cape Town metros versus Mangaung metro and Capricorn and NM Molema DMs), they hypothesized a higher burden of stigma in smaller metros and DMs likely posed significant barriers to MSM linking to care; they endorsed two sets of extrapolation assumptions as the median indicator estimates of the 2 large metros [A], and the median of the three less urbanized [B and C] sites. We present selected extrapolated MSM cascade results for three municipalities (Tshwane metro with Category A assumptions; Buffalo City metro and Ehlanzeni district municipality with Category B assumptions) in Figure 3.

**Figure 3.**
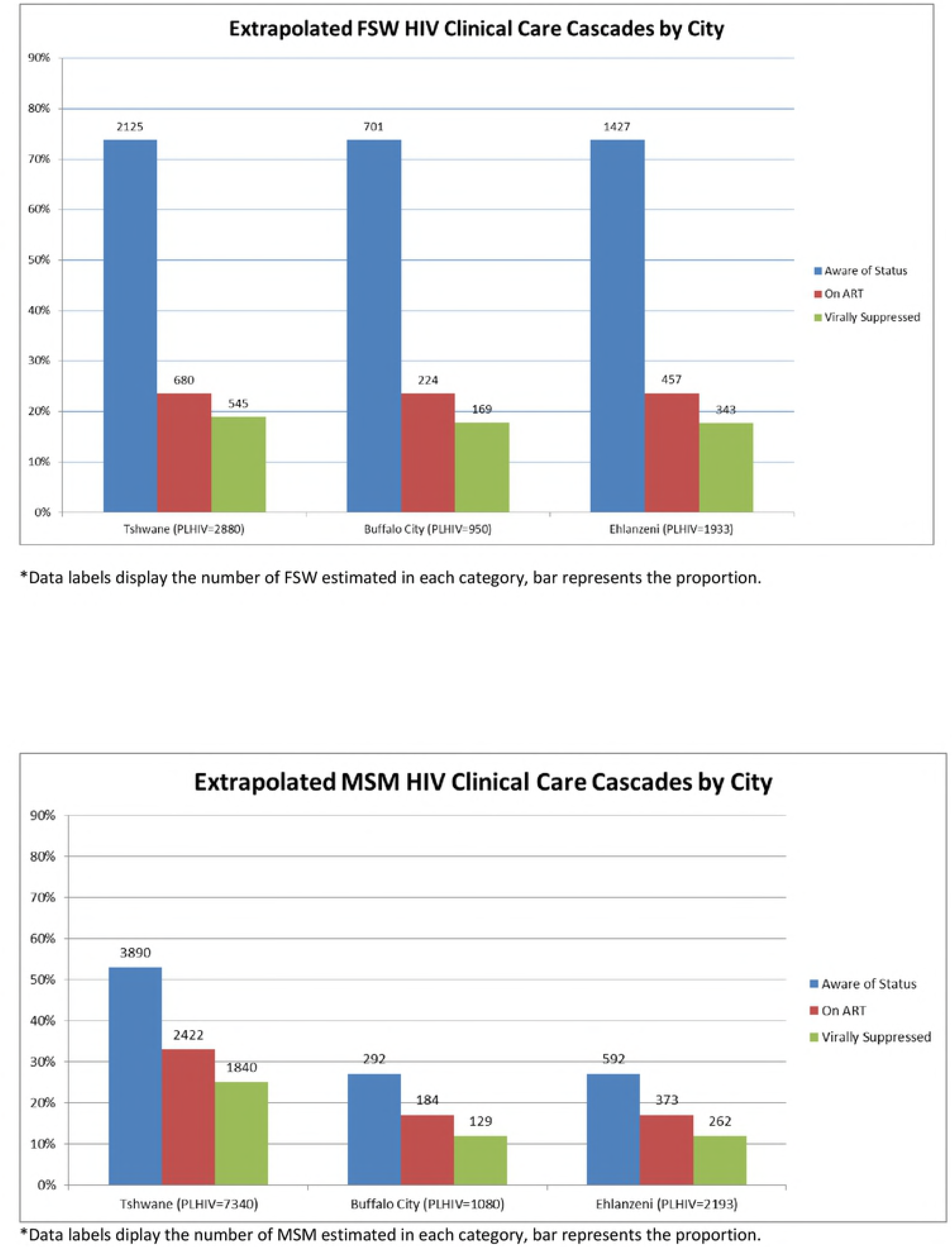
Selected Sub-National HIV Treatment Cascades for FSW and MSM (Extrapolated), 2018.

## Discussion

Our cascades analysis of South African FSW and MSM surveillance data visually demonstrates two population-specific trends in HIV testing and treatment access. Among FSW, we observed high awareness of HIV status but a steep breakpoint to ART uptake. By comparison, among MSM, we observed less awareness of HIV status, but relatively good ART uptake among those aware of their status. These results were consistent with stakeholders’ impressions from their own programmatic work, where FSW were generally receptive to testing but difficult to link to and retain in care, and MSM difficult to encourage to test but motivated to link to treatment upon diagnosis. It is important, however, not to overstate these broad comparisons into assumptions about any particular organization’s reach, or the effectiveness of any particular programmatic initiative. Apart from FSW awareness of status, which approached optimal levels only in Johannesburg, all other treatment cascade indicators were well below UNAIDS 90-90-90 targets. The data suggest clear disparities between metropolitan MSM residing in relatively more tolerant social environments who have access to a wider range of public and private health services (including “specialty” community-based clinical services promoted through targeted MSM outreach); versus those living in smaller urban or rural districts characterized by higher levels of social stigma and few MSM-competent primary health clinics. And by their nature, cascades are silent about whether the “unreached” differ in salient ways from the “reached” in general, or in any particular district. Overall these cascades indicate plenty of work remains to be done for FSW and MSM. Moreover, KP needs are often distinct between and within KP groups and contexts. At sub-national levels, these comparisons allow stakeholders to visualize gaps in service provision (e.g. FSW linkage and retention; MSM regular HIV testing) to target intensive investment in programmatic activity consistent with 90-90-90 goals.

Additionally, while our viral load extrapolation assumption (parity with women and men in the general population) may at first glance appear not to account for the considerable number of social-structural obstacles key populations face in accessing or adhering to treatment, the triangulation of surveillance data with stakeholder programmatic data and opinion suggested otherwise. It is true that both populations face significant stigma in South African society, and sex work remains criminalized. The stakeholders critically vetted the extrapolation assumptions and the preliminary results in light of these factors, but ultimately the assumptions were consistent with their own clinical monitoring data, which they shared with the stakeholder group. This gives us further confidence in our conclusion that much of what predicts poor HIV outcomes for key populations in South Africa occur upstream in the cascade; and here is where focused and specific programmatic attention is required. FSW are lost between diagnosis and retention, MSM avoid testing and thus diagnosis; but those who do overcome population-specific obstacles may have the same chance at living healthy lives with HIV as the general population. We offer this conclusion with caution, in hope that it may be validated in the future with surveillance laboratory data (and that it will never again require extrapolation!). This said, despite this being a limitation of our cascade analysis, it demonstrates another strength of our consensus method, as laboratory data to complete the cascade will be unavailable for each population until 2019 and 2020 respectively. We would not want readers to conclude that surveillance viral load testing of KP samples is not necessary. Rather, we recommend this type of data triangulation within a consensus process as a temporary way forward, until laboratory analysis of surveillance biological samples is adequately funded and stakeholders can be confident in the validity of results.

The consensus process allowed us to confront common pitfalls in constructing cascades: that relevant data are frequently end products of diverse sources, methodologies, and indicator definitions (36, 37). In our case, investigators proceeded from best practice surveillance methodologies, and the FSW and MSM surveys tended to define cascades-relevant indicators in the same way; yet laboratory methodologies differed between the FSW and MSM IBBSs. Our consensus methodology was designed to encompass the breadth of methods and definitions and to articulate a unified, transparent, and consistent cascades methodology that proceeded from specific data sets and allowed for standardized extrapolation into data-scarce sub-national districts. Also, as demonstrated by the adjustment of FSW PSE in February 2018, this participatory method enables the reconsideration of previous estimates when new data comes to light that stakeholders agree by consensus should be taken into consideration. Such consensus is critical to the development of a coordinated and coherent UTT programmatic response and assessment of progress towards 90-90-90 targets. In our case, the consensus process unfolded alongside multiple strategic planning processes including the development of the National Strategic Plan for HIV, TB and STIs (NSP) and specific Sex Worker and LGBTI HIV plans; the finalization of sub-national cascades with viral suppression estimates occurred alongside the finalization of the NSP’s companion Provincial and District Implementation Plans. Stakeholders may still have opinions about the precision or accuracy of any individual estimate in any district; but they proceed from a shared understanding of what surveillance data suggests about current programmatic reach and future needs.

We acknowledge several limitations of this process that were explicitly vetted by our stakeholder group in arriving at the consensus estimates presented here. First, this process depends on the existence of a critical mass of stakeholders and surveillance and program data. The “KP cascades” here are explicitly FSW and MSM cascades. An opportunity to present data from 2 PWID and 2 transgender women’s surveillance sites as cascades is forthcoming in 2019. This progress in KP strategic information still lags well behind need; we emphatically call for the investment of resources in PWID and transgender programming and strategic information commensurate with realizing and tracking progress towards 90-90-90 for them in South Africa and elsewhere. Moreover, these cascades are informed exclusively by surveillance data. We will be implementing a protocol in 2019 to incorporate routinely collected monitoring and evaluation data into all KP cascades using the same participatory triangulation methodology with stakeholders. We believe that this planned triangulation of surveillance and program data with stakeholders’ collective experiences will result in potentially more robust estimates. Although the consensus process is transparent about the limitations of surveillance methods and data, it does not, strictly speaking, account or correct for these limitations, and may reproduce known and unknown biases. For example, in a case where PSE methods or IBBS survey recruitment may have systematically produced a point estimate biased towards an overestimate for any site (e.g. a rural district), this result will inform extrapolation assumptions and reproduce overestimates for multiple similar sub-national units. In our case among MSM, groups like university students may have been overrepresented in some surveillance samples, and cascades could be biased towards an underrepresentation of less educated MSM who lack access to resources and MSM-friendly health services (note that RDS adjustment was not consistently possible with our MSM surveillance data). Moreover, programmatic reach may be poorly aligned with surveillance survey inclusion criteria. The FSW and MSM surveillance surveys included young adults aged 16-17 years, but KP programs may lack substantial reach into adolescent sub-populations. Misalignment may ultimately be of little consequence to data interpretation and program planning, but it must be acknowledged and explicitly vetted by stakeholders. And, although the process overall affords stakeholders the opportunity to evaluate the precision of extrapolated results in any sub-national unit against their own knowledge and experience, extrapolation assumptions may nonetheless be biased towards the KP data- and service-rich urban environments.

Finally, like any inherently political process, it is entirely possible for the group to reach conclusions that may serve the interests of just a few organizational stakeholders through systematic over- or under-estimation, which may implicitly affect perceptions of their programmatic performance. Therefore, among the most important role the technical advisors play in this process is emphasizing that a gap between empirical data, program data, and expert opinion does not mean that any stakeholder is underperforming, but that a plausible degree of uncertainty exists. This can decrease the likelihood that “consensus” does not over- or understate need or coverage related to any individual stakeholder’s performance. And we concede here that although stakeholder vetting encouraged lively debate at times from all participants, it is of course no guarantee that we arrived at the absolute truth. Nonetheless, the stakeholder group itself must be satisfied that their consensus is accurate *enough* for effective programmatic planning.

## Conclusion

KP cascades could be constructed by one or two data analysts in a room by themselves on a shared workstation; we highly recommend that they not be (see Figure 4). We acknowledge that the participatory and inclusive approach co-created by stakeholders, technical analysis, and civil society is labor and resource intensive, and observe that these investments have already achieved positive impacts in wider data dissemination for program planning in South Africa. We trust it can do so elsewhere in a region where KP are frequently and deliberately excluded from both process and programs. We concede that the legal and human rights context of South Africa may be exceptionally enabling for FSW and MSM stakeholders and advocates; in some contexts similar stakeholder meetings may be prohibited by law or policy designed to prevent or suppress exactly this type of stakeholder input and empowerment. We hope this analysis aids their efforts to change this dynamic in national HIV planning.

**Figure 4.**
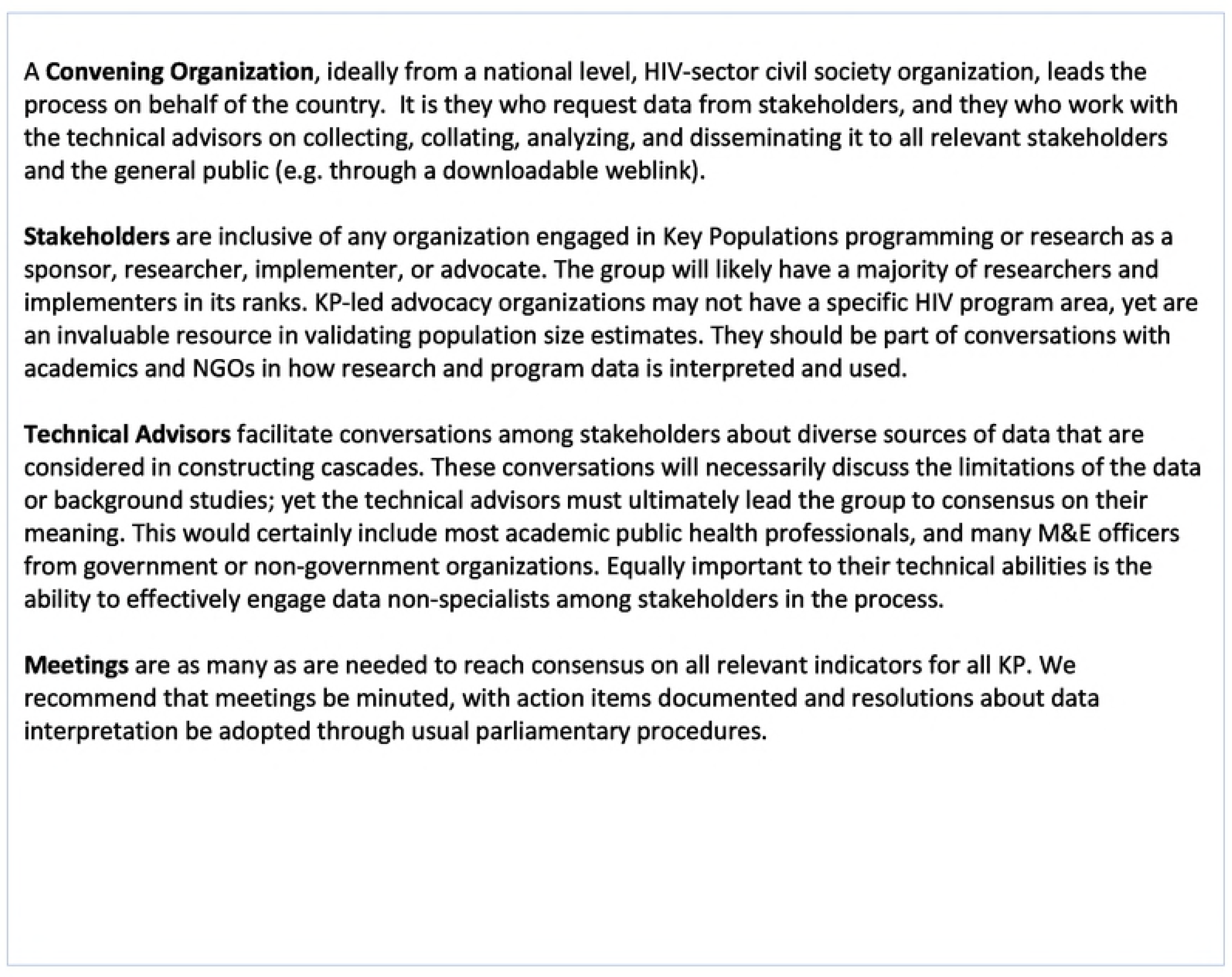
Recommendations for a participatory consensus process for KP cascades.

